# tMHG-Finder: Tree-guided Maximal Homologous Group Finder for Bacterial Genomes

**DOI:** 10.1101/2025.03.16.643543

**Authors:** Yongze Yin, Bryce Kille, Huw A. Ogilvie, Todd J. Treangen, Luay Nakhleh

## Abstract

A *maximal homologous group*, or MHG, as a group of sequences with a shared evolutionary ancestry, shifts the focus from a genecentric view to a homology-centric view in comparative genomic studies. Each MHG is formed by identifying and grouping all homologous sequences, which ensures that evolutionary events, such as horizontal gene transfer, gene duplication and loss, or *de novo* sequence evolution, are encapsulated within the same MHG. However, the current MHG computation tool, MHG-Finder, faces challenges in scalability to handle large datasets and lacks the ability to provide detailed insights into intermediate MHGs involving subsets of input genomes. We present tMHG-Finder (https://github.com/yongze-yin/tMHG-Finder), a new method that improves our previous method, MHG-Finder, by utilizing a guide tree to significantly improve scalability and provide more informative biological results. We also introduce a new measure, fractionalization (available at https://github.com/yongze-yin/Fract-Calculator), to assess the accuracy of delineated MHGs compared to ground truth data. Our results show that tMHG-Finder scales linearly with the number of taxa, requiring a small fraction of the computational time of MHG-Finder. Furthermore, according to the fractionalization measure, tMHG-Finder outperforms four state-of-the-art whole-genome aligners on simulated data. Applying tMHG-Finder to a phylum of extreme-environment-resistant bacteria, we validated our results through the encapsulation of 16S rRNA sequences within MHGs. We further investigated how evolutionary rates change with phylogenetic distance and explored the functional roles of genes captured by conserved MHGs, demonstrating the broader utility of tMHG-Finder in uncovering evolutionary insights beyond MHG delineation and phylogenetic relationships.

## 1 Introduction

Bacterial evolution is a complex process affected by frequent genetic events and adaptations to the fast-changing environment [23,12,32]. Homologous groups, each defined as a set of sequences that share a common evolutionary ancestry, are used as loci to understand the evolutionary process. Traditional approaches rely on using functional gene sets as homologous groups [21]. However, this violates the fundamental mathematical assumption that the evolutionary history of each locus should be recombination-free [34]. In contrast, alignment-based methods provide a viable option for studying bacterial evolution without relying on protein annotation. These methods use the inherent genomic similarity of homologous sequences. However, existing whole-genome aligners are challenged at deeper evolutionary timescales where genomes are highly divergent.

Whole-genome alignments play a crucial role in downstream analyses in comparative genomic studies [19]. Particularly, they can serve as the foundation for constructing pan-genomes [8], a field of increasing attention due to its ability to reveal the genetic diversity and key genomic components contributing to bacterial adaptation. Pan-genome studies delineate the core and dispensable genomes, offering insights into bacterial genetic diversity. However, limitations in existing pan-genome construction methods hinder their impact. For example, some methods require the identification and annotation of protein sequences rather than solely relying on genomic sequences as input [28,25,31].

A *maximal homologous group*, or MHG, is defined as a maximal set of maximum-length sequences whose evolutionary history is a single tree [34]. Similar to the property of a locally collinear block (LCB), which denotes a rearrangement-free homologous region, the definition of MHG imposes two additional key optimality constraints. The set is maximal in the sense that no sequences that are not in the set are homologous to those in the set without involving internal rearrangements. Each sequence within the group is also maximal, meaning that flanking base pairs on either side of any sequence in the group do not belong to the homology group without involving internal rearrangements. Our previous alignment-graph-based MHG detector, MHG-Finder [34], is one of the few methods available for identifying MHGs directly. However, it has scalability and information limitation challenges.

To overcome these challenges, we introduce tMHG-Finder, a *de novo* tree-guided MHG finder. tMHG-Finder considers the evolutionary distances between the input genomes to adjust the expected genomic similarity accordingly and computes MHGs for both the complete set and subsets of input genomes. To improve scalability, tMHG-Finder employs a progressive approach by constructing a representative sequence for each MHG at each node in the input tree. As a result, the number of pairwise blastn [2] comparisons drops from *O*(*n*^2^) to *O*(*n*), where *n* signifies the number of input genomes. Moreover, tMHG-Finder supports parallel processing by utilizing multiple threads to concurrently handle different connected components of the constructed alignment graph.

We assessed the accuracy of tMHG-Finder against four whole-genome aligners: Progressive Cactus, progressiveMauve, SibeliaZ, and Mugsy [4,11,22,3]. Considering each identified alignment as an MHG, we initially employed classical multiple sequence alignment (MSA) set comparison metrics: recall, precision, and F-score. Additionally, we introduced and applied a novel comparison metric, fractionalization. In contrast to existing metrics [13,6] that examine the alignment quality (gaps, mismatches), fractionalization compares the endpoints of the aligned regions to quantify how a reference alignment is fragmented into a query alignment set without considering the alignment quality. When the true alignment set serves as the reference set, fractionalization aids in measuring the quality of identified breakpoints for homologies included in the target alignment. On a set of simulated genomes, aligning MHGs identified by tMHG-Finder, we show that tMHG-Finder and Cactus produce genome alignments with an F-score above 0.90, whereas all other methods produce alignments with an F-score below 0.75. Comparatively, tMHG-Finder yields alignments with higher precision than Cactus, albeit with lower recall. However, alignments generated from tMHG-Finder-identified MHGs exhibit significantly less fragmentation than those identified by Cactus, showcasing superior accuracy.

To explore evolutionary dynamics, we applied tMHG-Finder to an extremophilic bacterial phylum: *Deinococcus-Thermus*. The accuracy of tMHG-Finder was validated by confirming that all 162 copies of near-complete 16S rRNA sequences from 80 studied genomes were encapsulated within a single MHG, indicating that tMHG-Finder effectively captures conserved genomic regions. To assess evolutionary rates, we quantified genomic distances using average pairwise average nucleotide identity (ANI) values. As genomes became more distantly related (lower ANI values), the number of MHGs increases, while the average length of MHGs decreased, consistent with expectations of increased divergence. By tracking MHG results across internal nodes of the guide tree, we observed a reduction in the average number of genes covered by MHGs as genomic distances increased. Focusing on conserved MHGs, defined as MHGs containing exactly one sequence from every studied genome, we conducted a gene ontology (GO) analysis and identified a diverse range of functional categories associated with the genes captured by these conserved MHGs, providing insight into the functional roles of genes under evolutionary constraints. These results highlight the relationship between evolutionary rates and phylogenetic distances, demonstrating the ability of tMHG-Finder to capture genomic evolution and offering insights into the evolutionary dynamics of extremophilic bacteria.

## 2 Materials and Methods

### tMHG-Finder Algorithm

Taking whole-genome nucleotide sequences as the sole input, tMHG-Finder partitions genomes into MHGs. In contrast to MHG-Finder presented in [34], tMHG-Finder utilizes a guide tree that can be either user-provided or estimated. As a result, tMHG-Finder employs a progressive approach by traversing the guide tree from the bottom leaf nodes to the top root node. At each internal node, genomes located at the leaves beneath the target internal node are partitioned into a set of MHGs, and then represented by a set of representative sequences constructed from the identified MHG set. Consequently, an upper-level internal node only needs to compute and partition the homologous matching pairs from the two sets of representative sequences of their direct children. This approach avoids the exhaustive all-versus-all blastn mappings and substantially reduces the blastn comparison complexity from *O*(*n*^2^) to *O*(*n*) where *n* is the number of input genomes at each internal node. Besides the enhanced computational efficiency compared to MHG-Finder, the final output of tMHG-Finder includes MHGs not only for the entire set of input genomes but also for the subsets along the evolutionary history.

To accurately infer a guide tree, tMHG-Finder leverages the MinHash-based alignment-free distance estimator Mash [24], enabling a fast calculation of pair-wise genomic distances between input genomes. Subsequently, neighbor-joining is employed on the distance matrix to sketch a guide tree determining the MHG partition order. Given a guide tree, the MHG partitioning at each internal node involves five steps: (1) blastn for pairwise local alignments, (2) sequence pile-up, (3) initial MHGs and alignment graph construction, (4) alignment graph traversal, and (5) representative sequence calculation. While the first four steps rely on the algorithm in [34], the final novel representative sequence calculation step results in substantial computational savings.

tMHG-Finder categorizes internal nodes into leaf internal nodes and non-leaf internal nodes. A leaf internal node has raw genome assemblies as both children, for which tMHG-Finder relies on MHG-Finder to compute MHGs. Notably, tMHG-Finder computes “representative sequences” for each MHG set to optimize computational efficiency. For single-sequence MHGs, indicating that they are not homologous with any leaf, these sequences are preserved within the parent node’s representative sequences for potential higher-level MHG partitioning. For each MHG containing multiple sequences, a representative sequence is derived through MSA using MAFFT [17] and majority voting for the most frequent nucleotide at each site, becoming part of the parent internal node’s representative sequence set. In contrast, for a non-leaf internal node for which at least one child is an internal node, it directly utilizes the two representative sequence sets of both children (or one representative sequence set and one raw assembly if a child is a leaf node) to search for pairwise local alignments and partition into MHGs. Each partitioning boundary on a representative sequence (representing a child MHG) is then mirrored back to genome sequences within that MHG. The final representative sequence calculation step eliminates the need for all-versus-all blastn alignments between leaf node genomes, resulting in tMHG-Finder’s capability to process a larger input scale.

To determine the homology threshold, tMHG-Finder adopts a dynamic parameter setting that adjusts to the evolutionary distance between genomes. Theoretically, as two species diverge over an extended period of time, their genomes become less similar despite sharing a common ancestor. tMHG-Finder integrates an estimated distance matrix generated by Mash and selectively fine-tunes blastn parameters based on the ground truth of a simulated dataset generated with realistic parameters. Specifically, tMHG-Finder defaults to three-parameter configurations using Mash and blastn. The selection of a configuration is determined by the minimum Mash-estimated pairwise sequence similarity within a given genome set, and each configuration corresponds to a specific task using blastn. Following the guidelines from the blastn manual, if the minimum pairwise similarity within the given genome set exceeds 95%, blastn employs the megablast task due to the high similarity of sequences. For pairwise similarities falling between 65% and 95%, tMHG-Finder relies on blastn to perform dc-megablast to avoid excessively partitioning genomes into small MHGs or being overly strict and identifying too few regions. If the minimum similarity is below 65%, traditional blastn is performed for inter-species comparisons.

### Fractionalization: MSA Comparison Statistic

Fractionalization measures the level of diversity or heterogeneity, and we have reformulated it to quantify the level of congruence between two sets of multiple sequence alignments (MSAs). Our proposed fractionalization value, denoted as *f*, is computed using

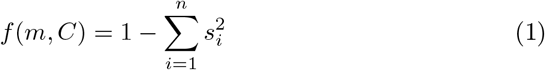

where *f* (*m, C*) represents the fractionalization value for a single MSA *m* distributed across *n* MSAs in the comparison set *C*. Specifically, *s*_*i*_ is the overlapping ratio, calculated as the sum of the lengths of overlapping ranges for sequences in *m* and *C*_*i*_, divided by the sum of the sequence lengths in *m*, providing a nuanced view by considering the extent of overlap. Taking two sets of MSAs as input, because fractionalization is an asymmetric metric, we treat each set alternately as the target and comparison set and compute the fractionalization value for every alignment within each set. The fractionalization value ranges from 0 to 1. As *f* approaches 0, it indicates high similarity, suggesting there exists an MSA in the comparison set *C* that closely matches the target alignment *m*. In contrast, as *f* approaches 1, it implies a higher degree of fragmentation, indicating that the target alignment *m* is fragmented into many different MSAs in *C*. The fractionalization value captures the spectrum of similarity and fragmentation of two sets of MSAs precisely and intuitively (Appendix Fig. S1).

### Simulation Study

We generated 30 bacterial genomes using the simulator ALF [10]. We first utilized ALF to build a birth-death model tree containing 30 extent leaves with a birth rate of 0.01 and a death rate of 0.001, and then evolved 30 genomes down the tree. At the root of the model tree, we initialized a genome with 4000 genes, approximately equal to the number of genes in an *E. coli* genome. The gene length was randomly drawn from a Gamma distribution with *k* = 3 and *θ* = 133.8. Evolutionary events were then introduced along the edges of the tree based on the probability of an event occurring on a single edge. The parameter settings were documented at https://github.com/yongze-yin/tMHG-Finder. Regarding coding versus non-coding length, we maintained a ratio of 85% to 15%, resembling the genomic composition of an *E. coli* genome. The coding regions were simulated using an M2 model, while the non-coding regions were simulated using a TN93 model. Additionally, the simulation provided the ground truth data including true loci, true gene trees, and the true species tree.

To benchmark the scalability and accuracy of tMHG-Finder in identifying MHGs, we compared tMHG-Finder v1.0.0 with four state-of-the-art wholegenome aligners: Cactus v2.3.0 [4], progressiveMauve with a build date of Feb 13, 2015 [11], SibeliaZ v1.2.4 [22], and Mugsy v1.2.3 [3]. Each alignment identified by these methods was regarded as an MHG, enabling a direct accuracy comparison against the ground truth. Progressive Cactus required a guide tree as input. To address this, we provided Cactus with the neighbor-joining topology tree (without branch lengths), which was estimated from the Mash distance matrix as part of the tMHG-Finder output. For SibeliaZ, we set *k*=15 as recommended by the SibeliaZ manual for bacterial datasets. The remaining parameters of each aligner were set to their default settings. The runtime benchmark was conducted using subsets of 30 simulated genomes, ranging from 2 taxa to the entire dataset. Eight threads were employed if a tool supported using multiple threads; otherwise, the tool was run with a single thread. Notably, when provided with a customized guide tree, tMHG-Finder allows users to obtain MHG sets for specific genome subsets in a single run, eliminating the need for multiple runs for different subsets.

### An Extremophilic Phylum of Bacteria

We investigated an extremophilic phylum of bacteria, *Deinococcus-Thermus* or *Deinococcota*, with tMHG-Finder. For each known species within the *Deinococcus-Thermus* phylum, we retrieved the highest-quality assembly from GTDB [26]. To ensure data quality and assembly completeness, We selected assemblies classified as GCF by NCBI and applied an additional filter using CheckM [27], retaining only those with completeness levels exceeding 99%. This curation process yielded a final dataset of 80 assemblies, comprising 52 species from *Deinococcus* and 28 from either *Thermus* or *Meiothermus*.

We applied tMHG-Finder to these 80 genomes and obtained intermediate MHG results for each internal node of the estimated guide tree. Each internal node on the guide tree is annotated with the number and average length of MHGs. To validate the accuracy of these results, we examined gene annotation files (GFF) obtained from NCBI to identify 16S rRNA locations annotated as “16S.” Given the highly conserved nature of 16S rRNA, its encapsulation within MHGs serves as a robust validation metric.

To quantify genomic distances, we utilized pyANI [29] which calculates pairwise ANI values using nucleotide BLAST (ANIb). Unlike alignment-free estimators such as fastANI [15], which struggle with ANI values below 80%, pyANI provides accurate results even for divergent genomes. The genomic distance for a given set of genomes was represented as the mean pairwise ANIb value.

We defined a “conserved MHG” as an MHG with exactly one sequence from each studied genome. For genes within conserved MHGs, we developed a pipeline to retrieve UniProt IDs and protein names [9] and query QuickGO [5] for gene ontology (GO) terms, using *Deinococcus radiodurans* (taxonomy ID: 243230) as the reference. Each retrieved GO term was propagated through the gene ontology tree to include all parent terms connected via “is_a” and “part_of” relationships. Functional categorization was achieved by intersecting the resulting GO terms with the Gene Ontology Prokaryote subset [1].

## 3 Results

### 3.1 tMHG-Finder Runtime Benchmark

To assess the scalability of tMHG-Finder in comparison to its predecessor and existing whole-genome aligners, we executed all tools using varying thread configurations on subsets of simulated data, gradually scaling up to the entire genome set. This experiment was conducted on an Intel Xeon Gold 5218 processor, utilizing eight threads if the tool supports multiple threads and one thread if not.

Compared to tMHG-Finder, MHG-Finder experienced a substantial increase in runtime when processing more than 6 genomes, taking 54 hours for 9 genomes. In contrast, tMHG-Finder processed 30 genomes in 13 hours using a single thread and 7 hours using eight threads (Fig. 2). Cactus and SibeliaZ took only 30 minutes to align 30 genomes, exhibiting high speed. Conversely, Mauve took 53 hours to process 30 genomes (Fig. 2).

**Fig. 1.**
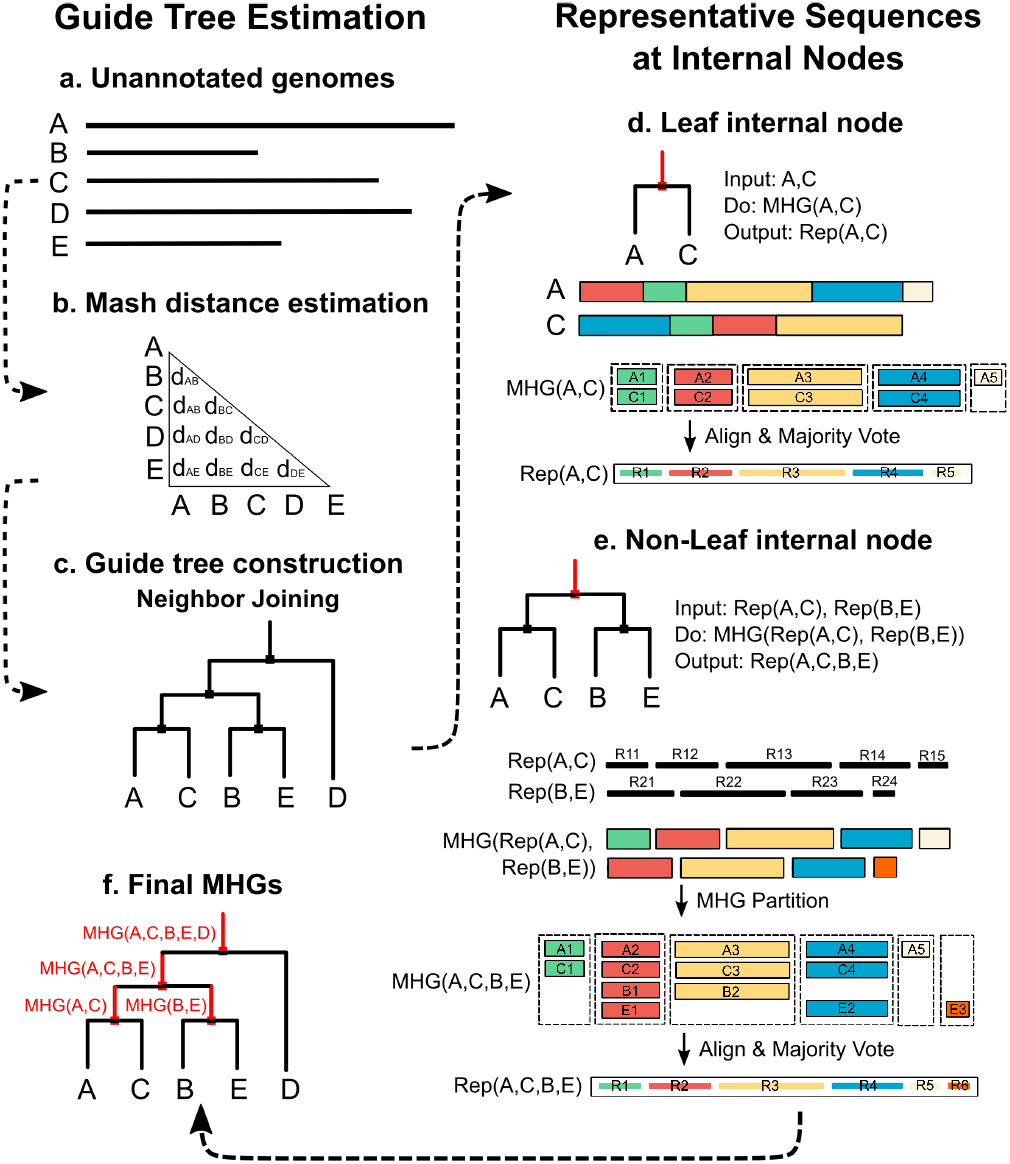
tMHG-Finder algorithm pipeline. (a): The input comprises a set of two or more genomes. (b): A distance matrix is estimated utilizing Mash. (c): A binary guide tree is constructed using the neighbor-joining method based on estimated pairwise distances by Mash. (d): A leaf internal node is an internal node with both children being leaf nodes. For every leaf internal node in the guide tree, tMHG-Finder utilizes the preceding MHG package MHG-Finder to compute MHGs. Upon obtaining a set of MHGs, tMHG-Finder computes a “representative sequence set” by aligning each MHG and employing majority voting to derive a consensus sequence representing the target MHG. (e): A non-leaf internal node is defined as an internal node with at least one child being an internal node. Instead of performing all-versus-all pairwise alignment for all included taxa under the target internal node, only the two children representative sequence sets are locally aligned and computed for MHGs. (f): The final output comprises an MHG set per internal node, which encompasses all taxa located beneath the respective internal node.

**Fig. 2.**
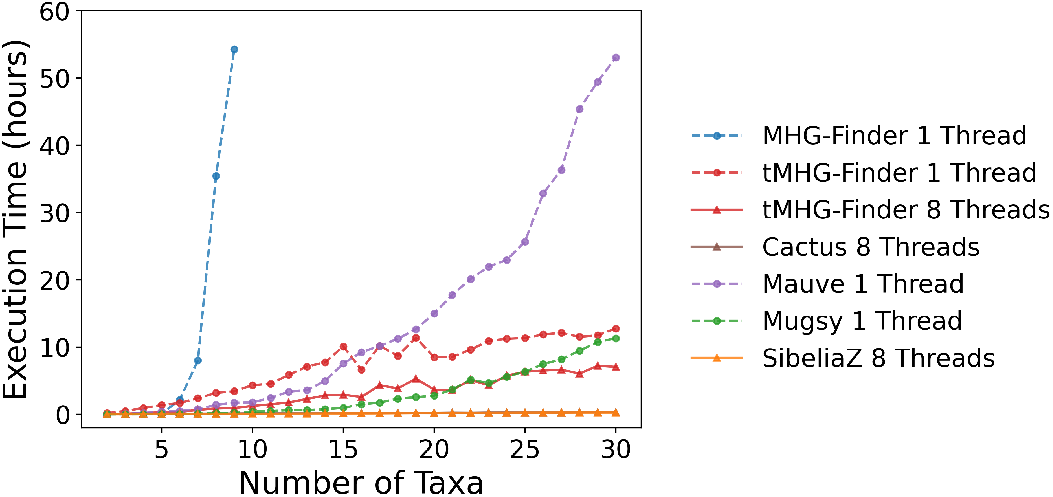
Execution time comparison. Wall time for computing MHGs using: MHG-Finder with one thread (blue), tMHG-Finder with one thread (red), tMHG-Finder with eight threads (red), Cactus with eight threads (brown), Mauve with one thread (purple), Mugsy with one thread (green), and SibeliaZ with eight threads (orange). Cactus and SibeliaZ overlap at the bottom. The execution time for Cactus does not include guide tree computation and hal-to-maf file conversion time.

### 3.2 tMHG-Finder Accuracy Assessment

To assess the accuracy of tMHG-Finder and existing whole-genome aligners at identifying homologous groups, we compared the MHG sets identified by each tool with the ground truth provided by ALF. The simulated dataset comprises 30 taxa and 3989 true MHGs. The number of sequences in each MHG roughly follows a normal distribution with a mean value of 30 (Appendix Fig. S2a). The length of each MHG, defined as the average sequence length, mostly ranges from 500 bp to 1,500 bp, with some extending close to 5,000 bp (Appendix Fig. S2b). The true pairwise Average Nucleotide Identity (ANI), calculated from the ground truth MSA, spans from 65% to 99%, with an average of 72% (Appendix Fig. S2c). To quantify the accuracy of each tool, we used mafComparator [13] to calculate the recall, precision, and F-score. mafComparator takes two sets of MSAs as input. For each set, it computes the ratio of the number of aligned nucleotide pairs in both sets to those in the target set. Recall quantifies the percentage of truly aligned pairs successfully identified by a tool, while precision represents the percentage of predicted aligned pairs present in the ground truth.

All tools demonstrated a precision over 97%, indicating their ability to identify aligned pairs correctly. While tMHG-Finder exhibited the highest precision, its recall was slightly lower than Cactus (Table 1). A noticeable observation was that SibeliaZ under-performed in terms of recall. We investigated and found that, unlike other tools with over 95% coverage for all genomes, SibeliaZ’s coverage is close to 50% for nine genomes, leading to a suboptimal benchmark for this metric. This is likely due to the high level of divergence and the fact that a 15-mer has *<*1% chance of remaining unmodified at a distance of 72% ANI [7].

**Table 1.** Recall, precision, and F-score comparison. Statistics computed using mafComparator comparing the simulated true MHG set with MHG sets identified by Cactus, tMHG-Finder (aligned by MAFFT v7.490), Mauve, Mugsy, and SibeliaZ.

| Tool | Recall | Precision | F-score |
| --- | --- | --- | --- |
| Cactus | **0.945** | 0.987 | **0.966** |
| tMHG-Finder | 0.877 | **0.994** | 0.932 |
| Mauve | 0.561 | 0.991 | 0.716 |
| Mugsy | 0.213 | 0.984 | 0.350 |
| SibeliaZ | 0.082 | 0.973 | 0.151 |

Additionally, we utilized fractionalization values to compare each identified MHG set with the true MHG set. As fractionalization is not symmetric, we computed this value by alternately considering each identified MHG set and the true MHG set as the target set and comparison set. This approach provides insights into whether a tool tends to over-fragment or under-fragment the truth.

SibeliaZ (Fig. 3: e and f) and Progressive Cactus (Fig. 3: g and h) exhibit a similar pattern, tending to identify short MHGs. Consequently, most true MHGs have a fractionalization value close to 1, indicating that true MHGs are fragmented into too many identified MHGs. SibeliaZ and Progressive Cactus are overly conservative, even when each identified group is truly homologous, though not necessarily a maximal group.

**Fig. 3.**
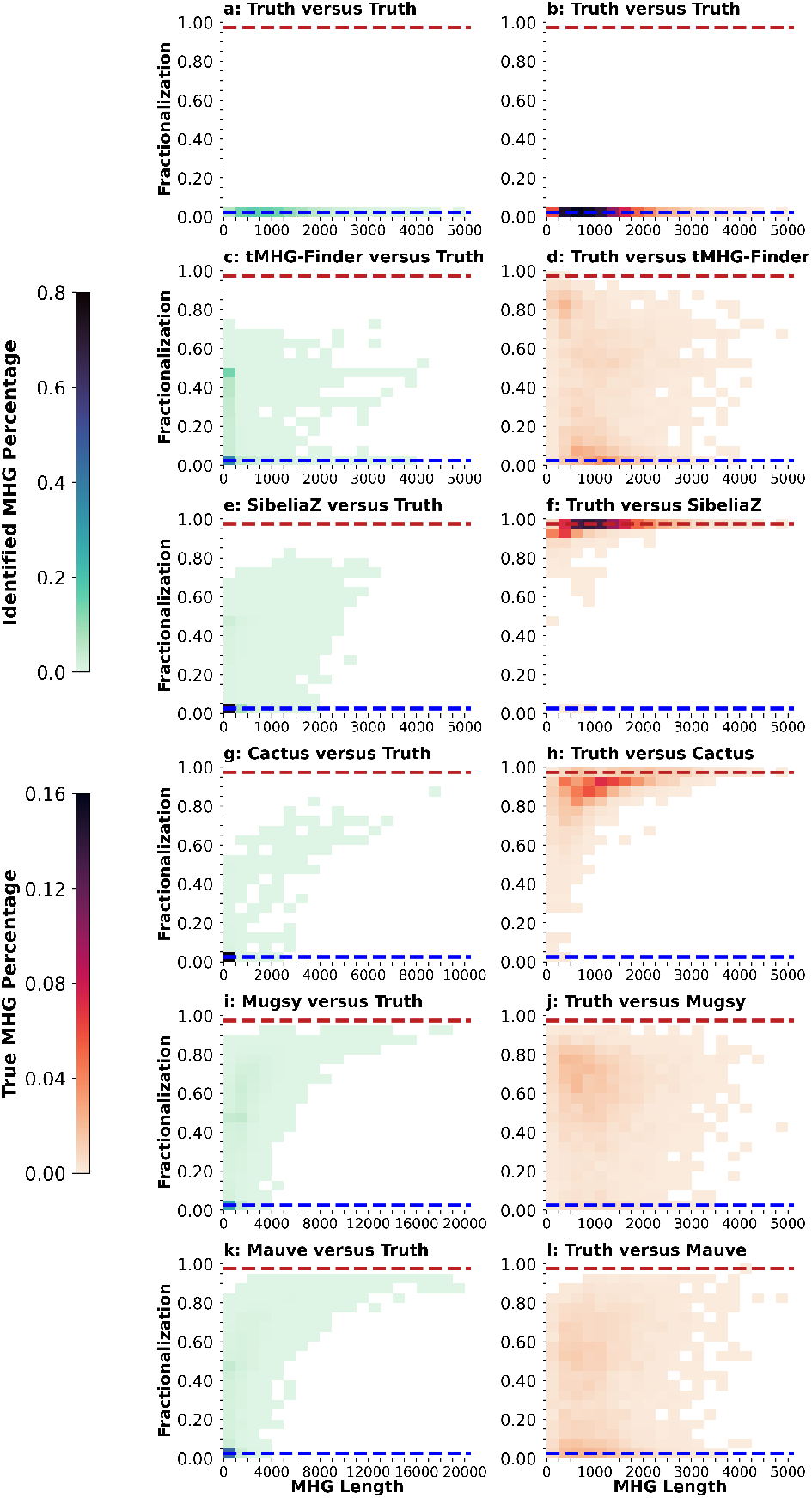
Benchmarking identified MHG accuracy with fractionalization. Each heatmap pair illustrates the distribution of fractionalization values: the left plot demon-strates how identified MHGs are fragmented into true MHGs, while the right plot demonstrates how true MHGs are fragmented into identified MHGs. Proximity to the red line in the left plot implies that the tool under-partitions the genomes, while proximity to the red line in the right plot suggests over-partitioning. Proximity to the blue line in both plots signifies high accuracy. Subplots (a, b): ALF Truth. (c, d): tMHG-Finder. (e, f): SibeliaZ. (g, h): Cactus. (i, j): Mugsy. (k, l): Mauve.

On the other hand, Mugsy (Fig. 3: i and j) and progressiveMauve (Fig. 3: k and l) identify MHGs ranging up to roughly 20,000 base pairs. As the length of an identified MHG increases, it naturally encompasses a larger number of true MHGs within it. Consequently, this leads to higher fractionalization values for identified MHGs. Notably, both algorithms demonstrate a relaxed approach, displaying a tendency to under-partition MHGs.

tMHG-Finder identifies MHGs ranging up to 4,500 base pairs, aligning closely with the truth. Although tMHG-Finder identified MHGs have some fractionalization values close to 0.5, the performance remains relatively stable as the MHG length increases. Whether considered as the target set or the comparison set, tMHG-Finder identifies MHGs that are closer to the truth, as evidenced by a majority of fractionalization values accumulating around 0 from both perspectives (Fig. 3: c and d).

### 3.3 Validation in *Deinococcus-Thermus* Genes and Functions

To validate the accuracy of tMHG-Finder, we analyzed 312 sequences annotated as “16S”, including both partial and complete 16S rRNA sequences [33,18], ranging from approximately 50 bp to 1,500 bp. Of these, 162 were near-complete (1,400 bp), while the remaining were mostly less than 200 bp. By intersecting the MHGs identified at the root node of the guide tree with the genomic coordinates of 16S sequences, we found that all 162 near-complete 16S sequences were encapsulated within a single MHG, with an average length of 1,256 bp. Shorter sequences were likely artifacts from sequencing or assembly errors, as 16S rRNA sequences are highly conserved, and short fragments often appear as isolated contigs. While the mathematical definition of an MHG assumes all observed homologous relationships result from true biological events, it does not account for such errors. To address this, tMHG-Finder employs heuristics to exclude homologous sequences that are too short, preventing excessive partitioning of MHGs into small units. This approach ensures MHGs reflect biologically meaningful homologous relationships, demonstrating the method’s ability to capture true patterns rather than artifacts.

Expanding the analysis beyond 16S rRNA, we examined the relationship between MHG coverage of genes and evolutionary distance. At each internal node of the guide tree, we identified overlaps between MHGs and annotated gene coordinates, defining a gene as “covered” if at least 75% of its length was encompassed by an MHG sequence. To quantify gene coverage, we normalized the number of covered genes by the number of sequences in the MHG, calculating the average gene coverage per MHG sequence.

From leaf nodes containing two genomes to the root node encompassing all 80 genomes, the average number of genes covered per MHG decreased from 4.34 to 0.95. This trend aligns with the observation in Fig. 4 that the average length of MHG decreases from approximately 8,000 bp in the leaf nodes to 1,000 bp at the root. At shallow evolutionary distances, MHGs often span multiple genes, exceeding gene boundaries. As the number of genomes increases and evolutionary distances grow, MHGs become shorter, often covering only portions of genes, reflecting reduced homology across their full length.

**Fig. 4.**
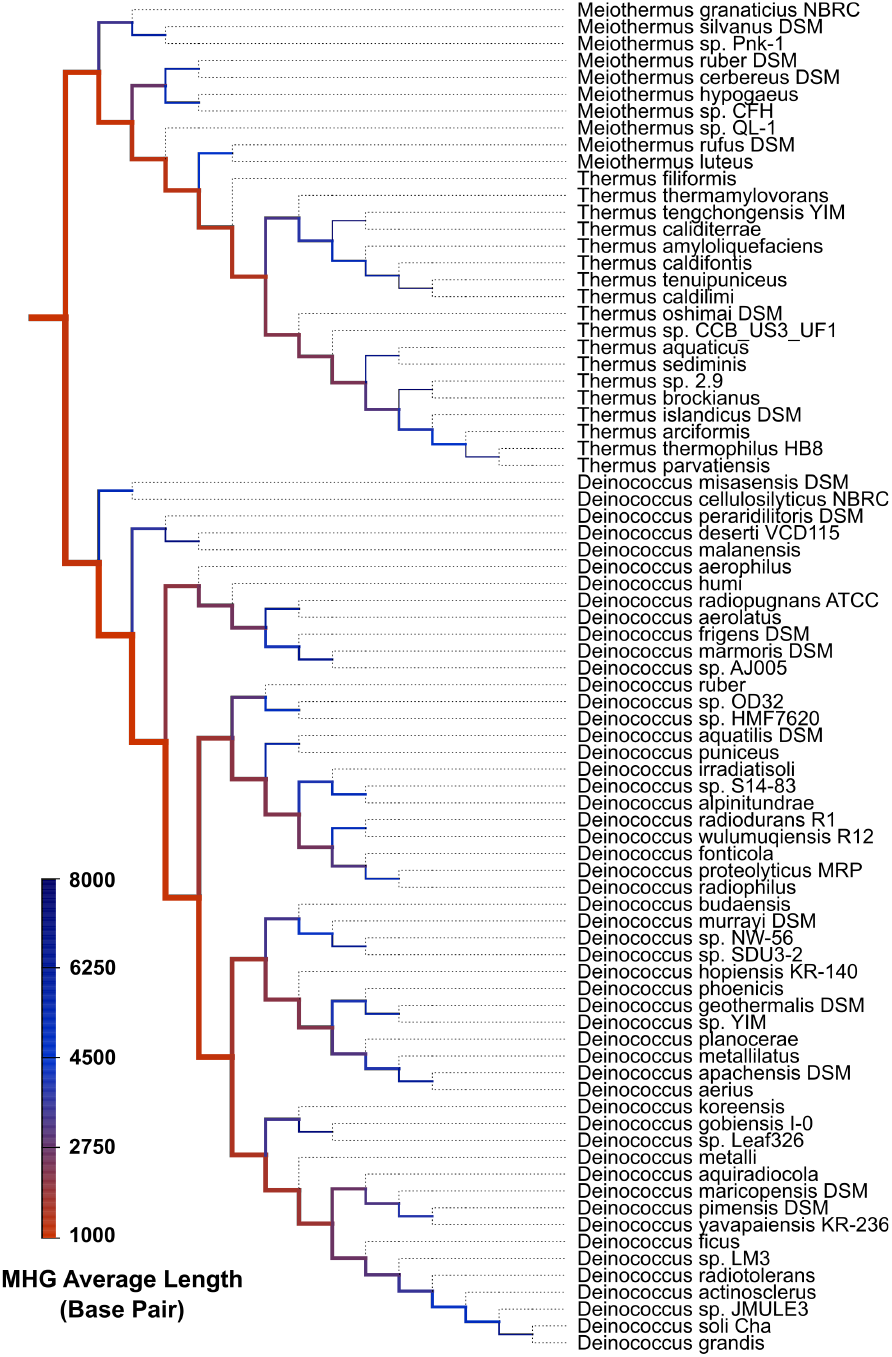
Annotated guide tree for the *Deinococcus-Thermus* phylum. Each tree branch is annotated with tMHG-Finder results corresponding to the linked internal node. The width of each branch indicates the number of MHGs that contain more than one sequence. A wider branch represents a higher count. The color of each branch, presented as a heatmap, reflects the average length of identified MHGs. The tip branches are represented as dotted lines, symbolizing the initiation of each genome as a single MHG with a length equivalent to the entire genome.

At the root node, we focused on genes covered by conserved MHGs. A total of 58 conserved MHGs were identified, with average sequence lengths ranging from 1,288 bp to 4,477 bp (Appendix Table S1). Of these, 14 MHGs overlapped with genes lacking annotated names. As expected, most conserved MHGs (38) corresponded to a single gene, with the longest being **rpoC**, which was fully encapsulated within its respective MHG. However, five MHGs spanned multiple genes. Among these, three contained multiple highly conserved ribosomal subunits, which are located next to each other and are conserved across all 80 genomes. The other two included **ClpX** and **ClpP**, involved in stress adaptation, and **sucC** and **sucD**, contributing to energy metabolism. An exception was observed with the gene **carB** splitting into two MHGs, each capturing approximately half of the gene. This is not a case of gene fission but rather an instance of over-fragmentation, highlighting a limitation of tMHG-Finder that may reduce its accuracy.

Conserved MHGs extended beyond ribosomal RNAs, prompting an analysis of their broader functional contributions. The genes covered by conserved MHGs were queried against the QuickGO database to retrieve their Gene Ontology (GO) terms, which were propagated through the ontology tree and intersected with the GO Prokaryote subset. While the most frequent terms were generic (e.g., MF: RNA binding, BP: translation, CC: cytoplasm), the GO terms contributed by MHG-covered genes spanned diverse categories (Fig. 5). These included functionalities potentially linked to the extremophilic traits of the studied phylum, such as DNA repair, various metabolic processes, and response to stimulus (Fig. 5). Investigating these functional contributions in greater detail remains a compelling future direction.

**Fig. 5.**
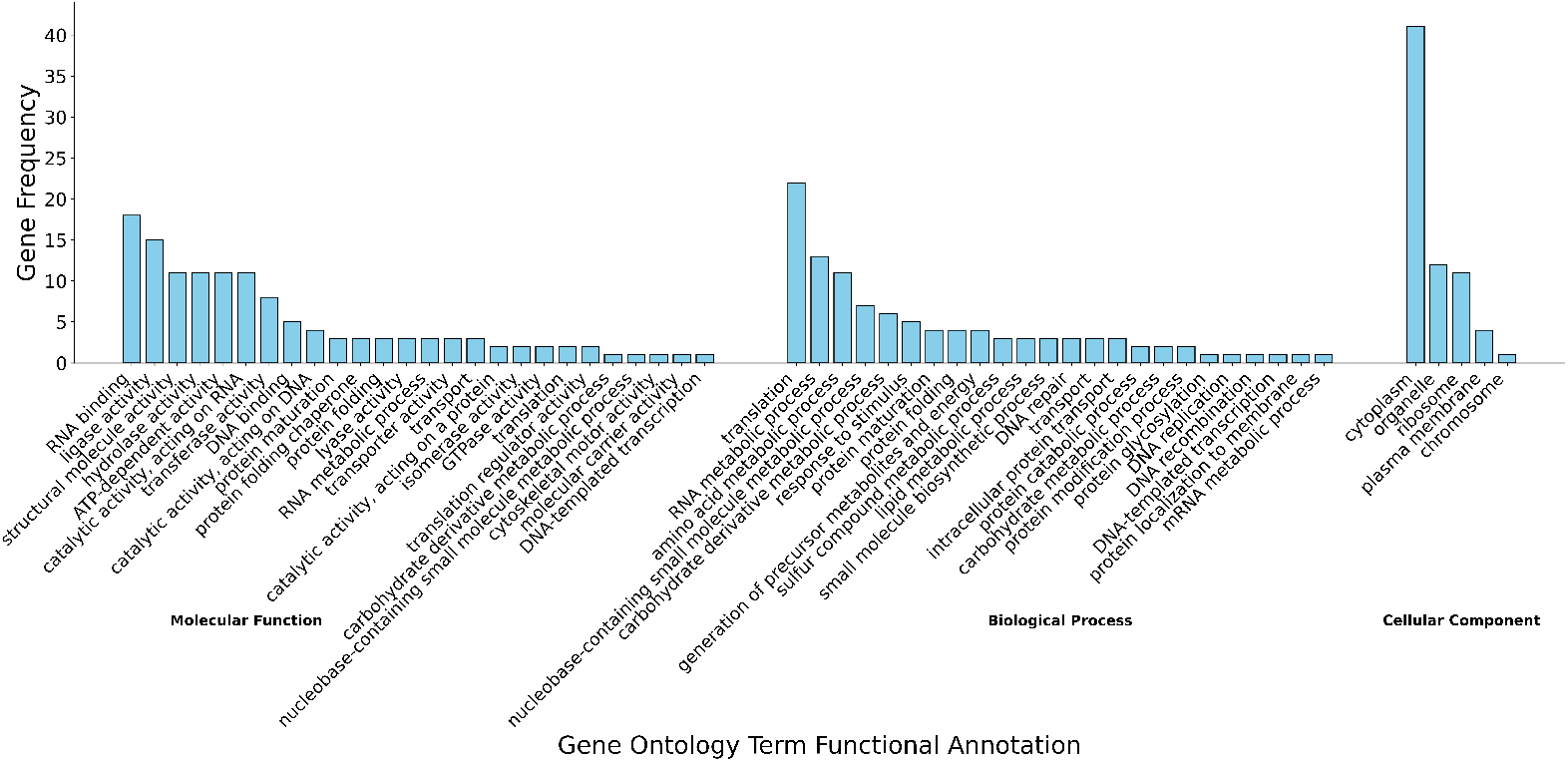
Gene Ontology Term Distribution Captured by Genes Covered by Conserved MHGs. The frequency distribution is categorized into three main activities: Molecular Function (MF), Biological Process (BP), and Cellular Component (CC). It is based on 54 genes encompassed by 44 conserved MHGs identified at the root node of the phylogenetic guide tree. For each gene, the associated Gene Ontology (GO) terms were propagated to include parent terms and intersected with the GO Slim Prokaryote subset to provide a high-level overview of functional contributions.

### 3.4 Deinococcus-Thermus Evolutionary Rates

In the analysis of the evolutionary rates of *Deinococcus-Thermus* phylum, the guide tree was constructed by merging two individual trees derived from 52 *Deinococcus* genomes and 28 *Thermus/Meiothermus* genomes, respectively. The guide tree was constructed using the neighbor-joining method based on the distance matrix estimated from Mash for the entire dataset. Since the dataset lacks an outgroup and the neighbor-joining tree is inherently unrooted, the tree was manually rerooted on the branch separating the two clades to provide a meaningful representation of the evolutionary relationships.

tMHG-Finder computed an MHG set at each internal node of the guide tree by including the genomes located at leaf nodes beneath that internal node. Each internal node was annotated with the number of MHGs with more than one sequence and the average length of identified MHGs. From the root to the leaves of the guide tree, there was a decrease in the number of MHGs and an increase in the average MHG length (Fig. 4). At the root node, including the most divergent species pairs within the studied dataset, 36,282 MHGs with an average length of 1,008 bp were identified. In contrast, internal nodes at the bottom level included greater conservation with less than 725 MHGs and a max length of 7,942 bp.

Zooming into specific clades (Fig. 4), the *Deinococcus* clade included 27,066 MHGs with an average length of 1048 bp. In contrast, the *Thermus/Meiothermus* clade, with 8,847 MHGs and an average length of 1010 bp, demonstrated a reduced number of MHGs compared to the *Deinococcus* clade. This observation was consistent with the fact that the *Thermus/Meiothermus* clade included about half the number of species found in the *Deinococcus* clade. Additionally, the average genome length of the *Thermus/Meiothermus* clade (2,599,819 bp) was substantially shorter than that of the *Deinococcus* clade (4,181,147 bp).

Quantifying genomic distance using the average ANIb across all genome pair combinations reveals a clear correlation with MHG statistics. As the genomic distance increases, the number of MHGs increases while their average length decreases. This trend is evident when comparing genome sets of similar size (Fig. 6 A and C). Another key factor influencing MHG statistics is the number of genomes in the compared sets. For example, B and C have similar average genomic distances, but B contains a larger number of genomes (indicated by higher transparency), exhibiting more MHGs with shorter lengths (Fig. 6). In summary, both including a larger number of genomes and increasing genomic distance lead to MHG sets containing a larger number of shorter MHGs.

**Fig. 6.**
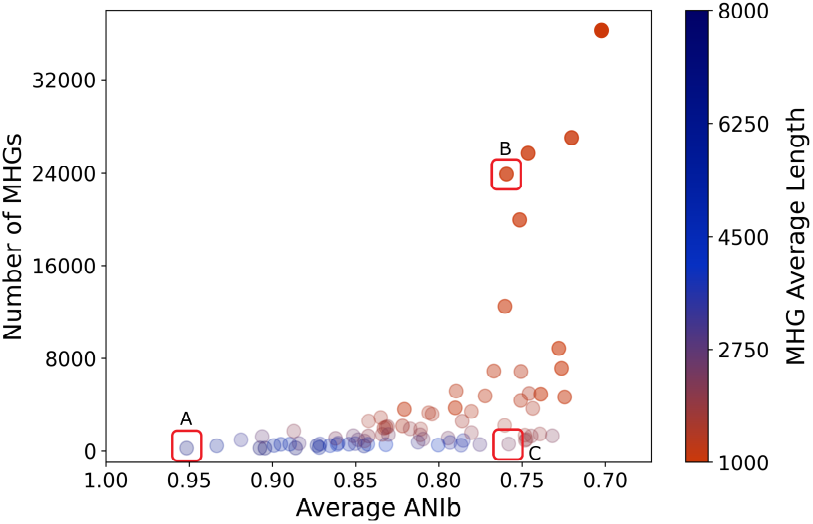
Correlation between ANI values, genome counts, and MHG statistics. Each dot represents an internal node on the guide tree. The x-axis shows the average ANI value between genome pairs in the left and right child nodes, while the y-axis indicates the number of non-singleton MHGs at the node. Dot color represents the average MHG length as a heatmap, and transparency reflects the number of genomes (ranging from 2 to 80). Points A, B, and C illustrate examples: A and C have similar genome counts but different ANI values, whereas B and C have similar ANI values but different genome counts.

The tMHG-Finder tree-based analysis demonstrated the complex relationship between evolutionary rates, phylogenetic distance, and homologous groups within the *Deinococcus-Thermus* phylum, providing valuable insights into the evolutionary processes of the extremophilic bacterial phylum.

## 4 Discussion

The definition of an MHG is rooted in an evolutionary perspective, imposing two crucial “maximal” constraints, ensuring that each MHG incorporates every homologous sequence and all sites for each sequence, sharing a single-tree history. These two constraints distinguish an MHG from a locally collinear block (LCB). It is crucial to note that MHG partitioning does not account for the order of neighboring MHGs. Additionally, MHGs allow for the inclusion of multiple sequences from the same genome, accommodating cases of duplication.

In this study, we present tMHG-Finder as an improvement to MHG-Finder. Leveraging a tree-guided architecture, tMHG-Finder dynamically adjusts the threshold to determine homologous sequences based on phylogenetic distances. tMHG-Finder accelerates blastn mapping with representative sequences and parallelization, enabling efficient processing of large datasets. It supports both user-defined and auto-estimated guide trees, as well as the integration of new genomes by comparing guide tree topologies and retaining MHGs below the insertion point. While guide trees and representative sequences are commonly used in genome alignment [4,11,30,16], they are seamlessly incorporated into tMHG-Finder. Unlike whole-genome alignment methods that prioritize maximizing pairwise alignment scores using the progressive framework, tMHG-Finder aims to optimize the evolutionary breakpoints at each internal node.

In contrast to classical clustering problems with well-defined elements in two compared clusterings, comparing two sets of MSAs in genome partitioning is challenging due to sequences rather than isolated elements in each MSA. Traditional metrics such as recall and precision, akin to the Rand index, treat each nucleotide as an individual element, overlooking that each sequence within an MSA represents a range. Fractionalization, as a novel metric, addresses these limitations by focusing on ranges and quantifying the congruence of breakpoints. It accurately captures the spectrum of similarity and fragmentation compared to existing metrics, providing a comprehensive assessment.

In our simulation study, our results show that tMHG-Finder is not only over 10x faster than MHG-Finder for less than 10 taxa on a single thread but also scales linearly with the number of taxa. Due to the limited scalability and less informative results of MHG-Finder, our evaluation focused on comparing the speed of the two versions of our tools. In contrast, for assessing accuracy, we focused on tMHG-Finder’s performance against four existing whole-genome aligners. Both tMHG-Finder and Cactus achieved F-scores above 0.90 (0.932 and 0.966, respectively), while the other methods’ F-scores were below 0.75. Cactus outperformed tMHG-Finder in recall (0.945 vs. 0.877), as tMHG-Finder adopts a more conservative approach in defining homologous group boundaries. Aiming to balance the two maximal constraints, the heuristics implemented in tMHG-Finder favor the maximal length constraint, which may result in lower recall but higher precision. However, Cactus exhibited a higher error rate, with 1.3% of its predicted alignments being incorrect, compared to only 0.6% for tMHG-Finder. This is noteworthy as tMHG-Finder’s predicted MHGs are significantly less fragmented than those from Cactus. The higher error rate of Cactus can introduce inaccuracies in downstream evolutionary analysis, which is why conservative core-genome alignment methods prioritize precision over recall [20,14].

The validation of tMHG-Finder through the accurate encapsulation of 16S rRNA sequences demonstrates the algorithm’s robustness. All near-complete 16S sequences were grouped into a single MHG, underscoring its capability to capture biologically meaningful homologous relationships while minimizing the impact of potential sequencing or assembly artifacts. Evolutionary rate analysis reveals a consistent trend: as species diverge, the number of MHGs decreases while their lengths increase, reflecting reduced conservation of homologous regions over evolutionary time. Additionally, the influence of genomic distance and dataset size on MHG statistics was evident. Larger datasets and greater genomic distances result in a higher number of shorter MHGs, while the average number of genes covered per MHG decreases with increasing evolutionary distance. This transition highlights the shift from MHGs spanning multiple genes at shallow evolutionary distances to covering portions of genes at deeper divergences.

The analysis of conserved MHGs further supports these observations, revealing that most overlap with single genes. These include ribosomal subunits and genes associated with key biological functions, such as stress response and energy metabolism. Notably, conserved MHGs within the *Deinococcus-Thermus* phylum capture genes contributing to extremophilic adaptations, including DNA repair, metabolic processes, and responses to environmental stimuli. These findings illustrate the ability of tMHG-Finder to identify functionally and evolutionarily significant homologous regions across diverse bacterial lineages.

In conclusion, tMHG-Finder provides a robust framework for investigating evolutionary dynamics through MHGs, offering valuable insights into bacterial adaptation to diverse environments. Future research could examine the influence of guide tree selection on MHG computation and assess the applicability of whole-genome aligners for comparative analysis within the *Deinococcus-Thermus* phylum. Additionally, exploring the evolution of key genes and operons through the lens of MHGs may further shed light on extremophilic traits and other adaptive mechanisms shaping bacterial evolution.

## Acknowledgments

This research was supported in part by the National Science Foundation, grants DMS/NIGMS-2153704, DBI-2030604, EF-2126387, and IIS-2239114.

## References

1. Aleksander, S.A., Balhoff, J., Carbon, S., Cherry, J.M., Drabkin, H.J., Ebert, D., Feuermann, M., Gaudet, P., Harris, N.L., et al.: The gene ontology knowledgebase in 2023. Genetics 224(1), iyad031 (2023)

2. Altschul, S.F., Gish, W., Miller, W., Myers, E.W., Lipman, D.J.: Basic local alignment search tool. Journal of molecular biology 215(3), 403–410 (1990)

3. Angiuoli, S.V., Salzberg, S.L.: Mugsy: fast multiple alignment of closely related whole genomes. Bioinformatics 27(3), 334–342 (12 2010). 10.1093/bioinformatics/btq665, https://doi.org/10.1093/bioinformatics/btq665

4. Armstrong, J., Hickey, G., Diekhans, M., Fiddes, I.T., Novak, A.M., Deran, A., Fang, Q., Xie, D., Feng, S., Stiller, J., et al.: Progressive cactus is a multiple-genome aligner for the thousand-genome era. Nature 587(7833), 246–251 (2020)

5. Binns, D., Dimmer, E., Huntley, R., Barrell, D., O’donovan, C., Apweiler, R.: Quickgo: a web-based tool for gene ontology searching. Bioinformatics 25(22), 3045–3046 (2009)

6. Blackburne, B.P., Whelan, S.: Measuring the distance between multiple sequence alignments. Bioinformatics 28(4), 495–502 (2012)

7. Blanca, A., Harris, R.S., Koslicki, D., Medvedev, P.: The statistics of k-mers from a sequence undergoing a simple mutation process without spurious matches. Journal of Computational Biology 29(2), 155–168 (2022)

8. Colquhoun, R.M., Hall, M.B., Lima, L., Roberts, L.W., Malone, K.M., Hunt, M., Letcher, B., Hawkey, J., George, S., Pankhurst, L., et al.: Pandora: nucleotide-resolution bacterial pan-genomics with reference graphs. Genome biology 22, 1–30 (2021)

9. Coudert, E., Gehant, S., De Castro, E., Pozzato, M., Baratin, D., Neto, T., Sigrist, C.J., Redaschi, N., Bridge, A.: Annotation of biologically relevant ligands in uniprotkb using chebi. Bioinformatics 39(1), btac793 (2023)

10. Dalquen, D.A., Anisimova, M., Gonnet, G.H., Dessimoz, C.: Alf—a simulation framework for genome evolution. Molecular biology and evolution 29(4), 1115– 1123 (2012)

11. Darling, A.E., Mau, B., Perna, N.T.: progressivemauve: multiple genome alignment with gene gain, loss and rearrangement. PloS one 5(6), e11147 (2010)

12. Darling, A.E., Miklós, I., Ragan, M.A.: Dynamics of genome rearrangement in bacterial populations. PLoS genetics 4(7), e1000128 (2008)

13. Earl, D., Nguyen, N., Hickey, G., Harris, R.S., Fitzgerald, S., Beal, K., Seledtsov, I., Molodtsov, V., Raney, B.J., Clawson, H., et al.: Alignathon: a competitive assessment of whole-genome alignment methods. Genome research 24(12), 2077–2089 (2014)

14. Fruzangohar, M., Moolhuijzen, P., Bakaj, N., Taylor, J.: Coredetector: a flexible and efficient program for core-genome alignment of evolutionary diverse genomes. Bioinformatics 39(11), btad628 (2023)

15. Jain, C., Rodriguez-R, L.M., Phillippy, A.M., Konstantinidis, K.T., Aluru, S.: High throughput ani analysis of 90k prokaryotic genomes reveals clear species boundaries. Nature communications 9(1), 5114 (2018)

16. Kaduk, M., Sonnhammer, E.: Improved orthology inference with hieranoid 2. Bioinformatics 33(8), 1154–1159 (2017)

17. Katoh, K., Standley, D.M.: Mafft multiple sequence alignment software version 7: improvements in performance and usability. Molecular biology and evolution 30(4), 772–780 (2013)

18. Kembel, S.W., Wu, M., Eisen, J.A., Green, J.L.: Incorporating 16s gene copy number information improves estimates of microbial diversity and abundance. PLoS computational biology 8(10), e1002743 (2012)

19. Kille, B., Balaji, A., Sedlazeck, F.J., Nute, M., Treangen, T.J.: Multiple genome alignment in the telomere-to-telomere assembly era. Genome Biology 23(1), 182 (2022)

20. Kille, B., Nute, M.G., Huang, V., Kim, E., Phillippy, A.M., Treangen, T.J.: Parsnp 2.0: Scalable Core-Genome alignment for massive microbial datasets. Bioinformatics p. btae311 (05 2024). 10.1093/bioinformatics/btae311, https://doi.org/10.1093/bioinformatics/btae311

21. Lerat, E., Daubin, V., Ochman, H., Moran, N.A.: Evolutionary origins of genomic repertoires in bacteria. PLoS Biology 3(5) (Apr 2005). 10.1371/journal.pbio.0030130

22. Minkin, I., Medvedev, P.: Scalable multiple whole-genome alignment and locally collinear block construction with sibeliaz. Nature communications 11(1), 6327 (2020)

23. Nakhleh, L., Ruths, D., Wang, L.S.: Riata-hgt: a fast and accurate heuristic for reconstructing horizontal gene transfer. In: Computing and Combinatorics: 11th Annual International Conference, COCOON 2005 Kunming, China, August 16–19, 2005 Proceedings 11. pp. 84–93. Springer (2005)

24. Ondov, B.D., Treangen, T.J., Melsted, P., Mallonee, A.B., Bergman, N.H., Koren, S., Phillippy, A.M.: Mash: fast genome and metagenome distance estimation using minhash. Genome biology 17(1), 1–14 (2016)

25. Page, A.J., Cummins, C.A., Hunt, M., Wong, V.K., Reuter, S., Holden, M.T., Fookes, M., Falush, D., Keane, J.A., Parkhill, J.: Roary: rapid large-scale prokaryote pan genome analysis. Bioinformatics 31(22), 3691–3693 (2015)

26. Parks, D.H., Chuvochina, M., Rinke, C., Mussig, A.J., Chaumeil, P.A., Hugenholtz, P.: Gtdb: an ongoing census of bacterial and archaeal diversity through a phylogenetically consistent, rank normalized and complete genome-based taxonomy. Nucleic acids research 50(D1), D785–D794 (2022)

27. Parks, D.H., Imelfort, M., Skennerton, C.T., Hugenholtz, P., Tyson, G.W.: Checkm: assessing the quality of microbial genomes recovered from isolates, single cells, and metagenomes. Genome research 25(7), 1043–1055 (2015)

28. Perrin, A., Rocha, E.P.: Panacota: a modular tool for massive microbial comparative genomics. NAR genomics and bioinformatics 3(1), lqaa106 (2021)

29. Pritchard, L., Glover, R.H., Humphris, S., Elphinstone, J.G., Toth, I.K.: Genomics and taxonomy in diagnostics for food security: soft-rotting enterobacterial plant pathogens. Analytical methods 8(1), 12–24 (2016)

30. Schreiber, F., Sonnhammer, E.L.: Hieranoid: hierarchical orthology inference. Journal of molecular biology 425(11), 2072–2081 (2013)

31. Tonkin-Hill, G., MacAlasdair, N., Ruis, C., Weimann, A., Horesh, G., Lees, J.A., Gladstone, R.A., Lo, S., Beaudoin, C., Floto, R.A., et al.: Producing polished prokaryotic pangenomes with the panaroo pipeline. Genome biology 21, 1–21 (2020)

32. Treangen, T.J., Rocha, E.P.: Horizontal transfer, not duplication, drives the expansion of protein families in prokaryotes. PLoS genetics 7(1), e1001284 (2011)

33. Větrovský, T., Baldrian, P.: The variability of the 16s rrna gene in bacterial genomes and its consequences for bacterial community analyses. PloS one 8(2), e57923 (2013)

34. Yin, Y., Ogilvie, H.A., Nakhleh, L.: Annotation-free delineation of prokaryotic homology groups. PLOS Computational Biology 18(6), e1010216 (2022)

