## Supplementary Material for "tMHG-Finder: Tree-guided Maximal Homologous Group Finder for Bacterial Genomes"

### Appendix A: Supplementary Material

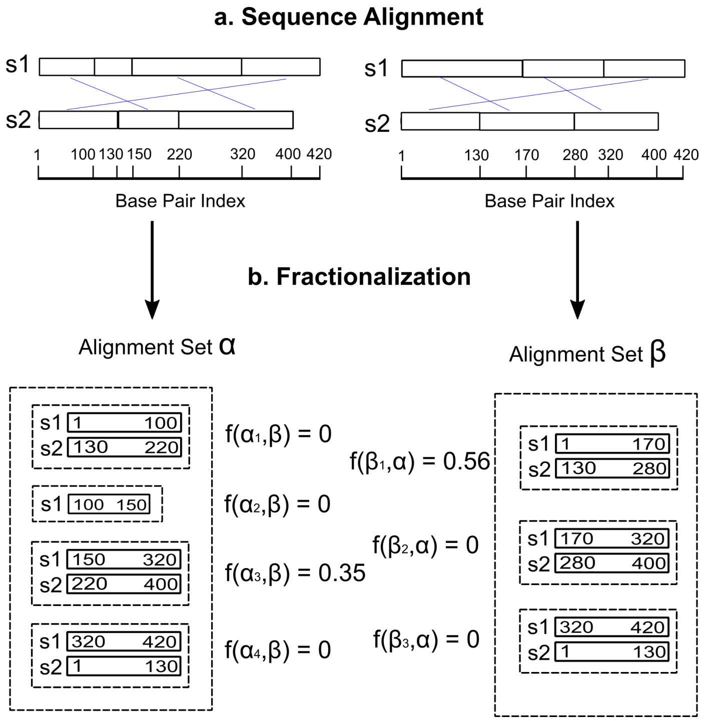


**Fig. S1. Quantifying congruence with fractionalization.** (a): Example sequences $s1$ and $s2$ are aligned in two ways, forming sets of alignments $\alpha$ and $\beta$. The blue line connects subsequences in the same alignment. (b): Comparing alignment sets $\alpha$ and $\beta$ using fractionalization (Eq. **??**). The value ranges from 0 (congruence) to 1 (fragmentation) for each alignment in each set, computed by alternatively considering two sets as the target set and the comparison set.

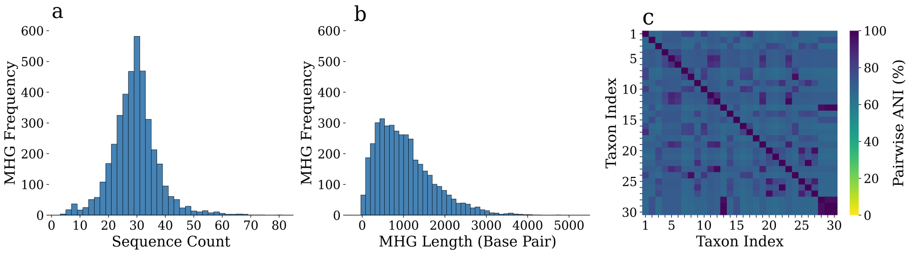


**Fig. S2. Simulation data characteristics.** (a): Distribution of the number of sequences within each MHG across the simulated dataset. (b): Distribution of MHG lengths in base pairs. (c): Average Nucleotide Identity (ANI) calculated from the true multiple sequence alignments for 30 simulated taxa.

Table S1: **Genes Covered by Conserved MHGs.** Gene names are obtained from NCBI gene annotations based on gene coordinates. UniProt IDs and protein names are queried from the UniProt database using the gene names. The length column represents the average length of the MHG sequences overlapping each gene.

| **Gene** | **Uniprot** | **Protein Name** | **Length** |
|---|---|---|---|
| accC | Q9RV16 | Acetyl-coenzyme A carboxylase carboxyl transferase subunit alpha | 1309 |
| acnA | Q9RTN7 | Aconitate hydratase A | 2637 |
| asnS | P0A8M0 | Asparagine–tRNA ligase | 1250 |
| aspS | Q9RVH4 | Aspartate–tRNA(Asp/Asn) ligase | 1587 |
| carB | Q9RWK0 | Carbamoyl phosphate synthase | 1313 |
| carB | Q9RWK0 | Carbamoyl phosphate synthase | 1480 |
| clpB | Q9RVI3 | Chaperone protein ClpB | 2156 |
| clpP | Q9RSZ7 | ATP-dependent protease proteolytic | 594 |
| clpX | Q9RSZ6 | ATP-dependent protease ATP-binding | 1141 |
| cysS | Q9RTT6 | Cysteine–tRNA ligase | 1237 |
| dnaK | Q9RY23 | Chaperone protein DnaK | 1865 |
| dxs | Q9RUB5 | 1-deoxy-D-xylulose-5-phosphate synthase | 1687 |
| fumC | Q9RR70 | Fumarate hydratase class II | 1329 |
| gatB | Q9RRD7 | Aspartyl/glutamyl-tRNA(Asn/Gln) amidotransferase subunit B | 1357 |
| glmS | Q9RXK9 | Glutamine–fructose-6-phosphate aminotransferase isomerizing | 1807 |
| gltX | Q9RX30 | Glutamate–tRNA ligase | 1345 |
| groL | Q9RWQ9 | Chaperonin GroEL | 1580 |
| guaA | Q9RT91 | GMP synthase glutamine-hydrolyzing | 1503 |
| gyrA | Q9RT53 | DNA gyrase subunit A | 1582 |
| ileS | Q9RUP8 | Isoleucine–tRNA ligase | 1269 |
| infB | Q9RTG5 | Translation initiation factor IF-2 | 1473 |
| lepA | Q9RV84 | Elongation factor 4 | 1765 |
| lysS | Q9RXE1 | Lysine–tRNA ligase | 1307 |
| malQ | P15977 | 4-alpha-glucanotransferase | 1299 |
| metG | Q9RUF3 | Methionine–tRNA ligase | 1286 |
| miaB | Q9RV79 | tRNA-2-methylthio-N(6)-dimethylallyl-adenosine synthase | 1259 |
| mnmG | Q9RYC3 | tRNA uridine 5-carboxymethylaminomethyl modification enzyme | 1742 |
| pnp | Q9RSR1 | Polyribonucleotide nucleotidyltransferase | 2050 |
| proS | Q9RUW4 | Proline–tRNA ligase | 1431 |
| purF | Q9RXT6 | Amidophosphoribosyltransferase | 1345 |
| purL | Q9RXT4 | Phosphoribosylformylglycinamidine synthase | 1524 |
| recG | Q9RT50 | ATP-dependent DNA helicase RecG | 1892 |
| rnr | Q9RZL8 | Protein NrdI | 1607 |

| **Gene** | **Uniprot** | **Protein Name** | **Length** |
|---|---|---|---|
| rplC | Q9RXK2 | Large ribosomal subunit uL3 | 625 |
| rplD | Q9RXK1 | Large ribosomal subunit uL4 | 618 |
| rplF | Q9RSL3 | Large ribosomal subunit uL6 | 529 |
| rplP | Q9RXJ5 | Large ribosomal subunit uL16 | 425 |
| rplR | Q9RSL2 | Large ribosomal subunit uL18 | 337 |
| rplV | Q9RXJ7 | Large ribosomal subunit uL22 | 309 |
| rplW | Q9RXK0 | Large ribosomal subunit uL23 | 279 |
| rpmC | Q9RXJ4 | Large ribosomal subunit uL29 | 161 |
| rpoC | Q9RVW0 | DNA-directed RNA polymerase | 4476 |
| rpsC | Q9RXJ6 | Small ribosomal subunit uS3 | 742 |
| rpsE | Q9RSL1 | Small ribosomal subunit uS5 | 422 |
| rpsJ | Q9RXK3 | Small ribosomal subunit uS10 | 317 |
| secA | Q9RWU0 | Protein translocase subunit | 1338 |
| secD | Q9RTE3 | Multifunctional fusion protein | 1611 |
| secY | Q9RSK8 | Protein translocase subunit | 1291 |
| speA | Q9RXR4 | Biosynthetic arginine decarboxylase | 1847 |
| sucC | Q9RUY3 | Succinate–CoA ligase (ADP-forming) | 1139 |
| sucD | Q9RUY2 | Succinate–CoA ligase (ADP-forming) | 862 |
| sufB | Q9RSL8 | SUF system FeS cluster assembly SufBD N-terminal domain-containing protein | 1262 |
| uvrA | Q46577 | UvrABC system protein A | 1296 |
| uvrB | Q9RS52 | UvrABC system protein B | 1797 |
